# Autonomous Dampening of Wnt Signaling Selects Zebrafish Spinal Cord Progenitors for an Oligodendrocyte Fate

**DOI:** 10.64898/2026.10.11.758343

**Authors:** Kimberly A. Arena, Santos J. Franco, Charles G. Sagerström, Bruce H. Appel

**Affiliations:** Section of Developmental Biology, Department of Pediatrics, University of Colorado Anschutz, Aurora, CO, 80045, USA

**Keywords:** OPC, specification, Wnt signaling, zebrafish, oligodendrocyte, glia

## Abstract

Nervous system development relies on coordinated formation of neurons and glia from neural progenitor cells (NPCs). During spinal cord development, NPCs in the pMN domain of the ventral spinal cord produce motor neurons followed by oligodendrocyte precursor cells (OPCs), a subset of which differentiate as oligodendrocytes, the myelinating cell of the vertebrate central nervous system. Whereas prior research has outlined an extensive gene regulatory network driving oligodendrocyte lineage cell differentiation, less is known about the mechanisms that specify NPCs for oligodendrocyte fate. To elucidate these mechanisms, our lab recently generated a single-cell RNA sequencing and single-cell ATAC sequencing multi-omics dataset from the zebrafish spinal cord. The dataset identified a molecularly distinct subset of NPCs that appear fated to the oligodendrocyte lineage, which we refer to as pre-OPCs. Our data revealed that pre-OPCs are enriched for RNAs encoding factors that antagonize Wnt signaling, such as *barhl2* and *cachd1*. The chromatin regions associated with these genes are highly accessible in pre-OPCs, suggesting that their expression is developmentally regulated. Using confocal light microscopy to visualize expression of a Wnt-responsive transgenic reporter, we found that Wnt signaling is downregulated in the spinal cord immediately prior to OPC specification. To explore how modulation of Wnt signaling influences OPC specification, we pharmacologically inhibited or activated Wnt signaling immediately prior to OPC specification. These experiments revealed that embryos treated with a Wnt inhibitor produced excess OPCs whereas those treated with a Wnt agonist had a deficit of OPCs. Additionally, CRISPR/Cas9 targeted mutations of *barhl2* and *cachd1* impaired OPC production. Finally, we identified *wnt4b* as a ventral source of Wnt and determined that loss of *wnt4b* enhances OPC specification. Together, our studies reveal pre-OPC modulation of ventral Wnt signaling as a mechanism of OPC specification in the spinal cord.

## INTRODUCTION

Establishment of functional neural circuits requires coordinated development of both neurons and glia. Neuronal axons are ensheathed by myelin, a specialized, lipid-rich membrane that is essential for proper action potential conductivity and supports neuronal health. In the central nervous system (CNS) of vertebrate animals, myelin is produced by glial cells known as oligodendrocytes. Understanding how oligodendrocytes are formed is essential to deciphering the development of functional neural circuits.

Neural progenitor cells (NPCs) located in the pMN domain of the ventral spinal cord, pMN progenitors, give rise to motor neurons, ventral interneurons, a subset of astrocytes, and ependymal cells (Floriddia et al., 2020; Masahira et al., 2006; Park et al., 2002; Zhou et al., 2001). Production of motor neurons and ventral interneurons begins as early as 10 hours post fertilization (hpf) in zebrafish (Park et al., 2004). Later in development, after neural tube formation, pMN progenitors begin producing oligodendrocyte precursor cells (OPCs) (Ravanelli et al., 2018; Ravanelli & Appel, 2015). Using fate-mapping and timelapse microscopy, we learned that motor neurons and OPCs arise from distinct pMN progenitors (Ravanelli & Appel, 2015). To determine the molecular identities of pMN progenitors fated for the oligodendrocyte lineage, we previously performed single-cell RNA-sequencing (scRNA-seq) of EGFP^+^ cells isolated from the trunks of *Tg(olig2:EGFP)* embryos (Scott et al., 2021). Using cell clustering and pseudotime trajectory analyses, we found a small population of transcriptionally distinct cells that appeared to represent a transition state between NPCs and OPCs, which we termed “pre-OPCs” (Scott et al., 2021). However, the molecular mechanisms that specify pre-OPCs for an oligodendrocyte fate remain unclear.

One signal transduction pathway implicated in OPC specification is the Wnt signaling pathway (Guo et al., 2015; Soomro et al., 2018). In the mouse cortex, Wnt signaling, revealed by a Wnt reporter gene, was high during neurogenesis but then much lower at the time of OPC production (Langseth et al., 2010), suggesting that Wnt inhibition might regulate OPC specification. Consistent with this, cortical expression of a dominant negative form of the Lef1 transcription factor, which blocks transcription of Wnt target genes, caused formation of excess OPCs (Langseth et al., 2010). By contrast, transgenic expression of *ß-catenin*, which activates Wnt signaling, under control of *Olig1* regulatory DNA impaired OPC formation in the mouse spinal cord (Dai et al., 2014). Together, these studies raise the possibility that a reduction of Wnt signaling in NPCs is necessary for them to adopt an OPC fate. However, whether Wnt signaling is dampened because of declining Wnt ligand availability or through NPC expression of genes that encode Wnt modulators is not known.

To dissect the regulatory mechanisms that modulate the OPC specification, we generated a scRNA-seq and single-cell ATAC-sequencing (scATAC-seq) multi-omics dataset (Arena et al., 2026). Using differential gene expression analysis, we discovered that pre-OPCs are uniquely enriched for RNAs encoding Wnt antagonists *barhl2* and *cachd1*. Chromatin is also highly accessible at these gene loci in pre-OPCs. This raised the possibility that pre-OPC expression of Wnt inhibitors selects these cells for oligodendrocyte fate. To investigate this possibility, we utilized in vivo pharmacological manipulation of Wnt signaling to explore its influence on OPC specification. Additionally, we used CRISPR/Cas9 mutagenesis to investigate the role of *barhl2* and *cachd1* on OPC generation. Finally, we applied single-cell multi-omics to identify *wnt4b* as a ventral source of Wnt and CRISPR/Cas9 gene editing to evaluate OPC production in zebrafish embryos lacking this ligand. Together, these studies present pre-OPC modulation of ventral Wnt signaling as a novel mechanism of OPC specification in the spinal cord.

## RESULTS

### pre-OPCs are enriched for transcripts that encode Wnt inhibitors and exhibit a distinct chromatin accessibility landscape

In prior work, we identified pre-OPCs as a transcriptionally distinct subset of neural progenitors that appear fated for the oligodendrocyte lineage (Scott et al., 2021). To uncover how these cells become transcriptionally distinct, we set out to identify genome-wide transcriptional and chromatin accessibility patterns unique to pre-OPCs. To do so, we generated a single-cell multi-omics dataset consisting of both single-cell RNA sequencing and single-cell ATAC sequencing. We sequenced cells sorted from embryos carrying a *Tg(olig2:egfp)* transgene, which labels pMN spinal cord progenitors and their progeny with EGFP, at 36 hpf when OPCs are specified (Arena et al., 2026). Cluster analysis of our multi-omics data revealed cell populations like those identified in our prior single-cell RNA sequencing data, including pre-OPCs (Scott et al., 2021)(Figure 1A). We reasoned that genes important for OPC specification would be expressed at higher levels in pre-OPCs than in other cell populations. We conducted differential gene expression analysis comparing pre-OPCs and neural progenitors (Figure 1B) and pre-MNs (Figure 1C). Relative to neural progenitors, pre-OPCs expressed RNA transcripts associated with an oligodendrocyte fate, such as *gsx2*, *notch3,* and *sox5* (Arena et al., 2026; Baroti et al., 2016; Xiao et al., 2019). When compared to pre-OPCs, pre-MNs expressed RNA transcripts associated with a neuronal fate, such as *isl1, elavl3,* and *slit3* (Carr et al., 2017; Costantino et al., 2024; Liang et al., 2011). Agreeing with our prior results (Scott et al., 2021), pre-OPCs are transcriptionally distinct from neural progenitors and the motor neuron lineage. Furthermore, the chromatin organization of pre-OPCs is distinct from neural progenitors and pre-MNs.

**Figure 1:**
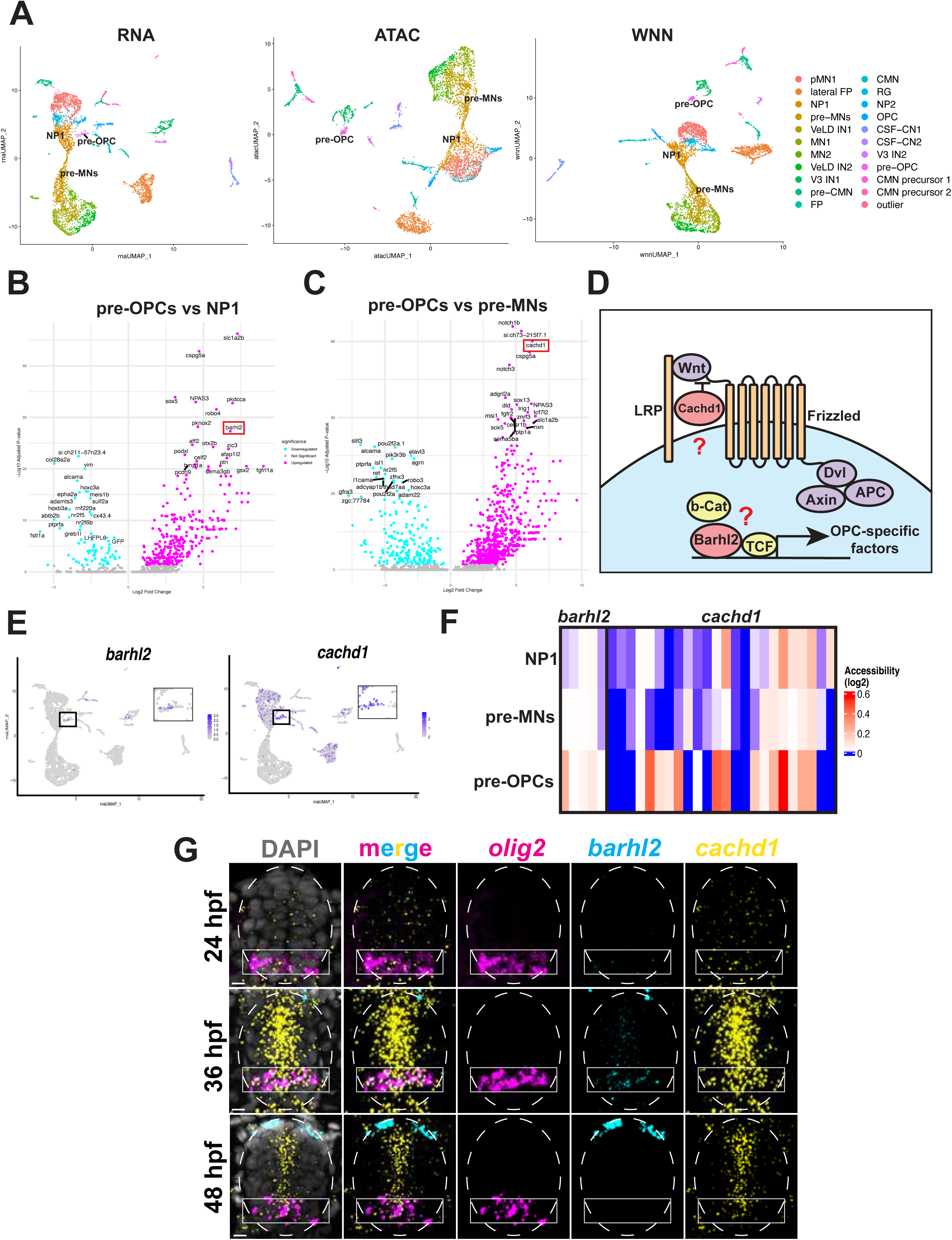
pre-OPCs are enriched for transcripts encoding Wnt inhibitors and exhibit a distinct chromatin accessibility landscape. A) RNA UMAP, ATAC UMAP, and Weighted Nearest Neighbors (WNN) UMAP generated from our scRNA-seq and scATAC-seq multi-omic dataset. WNN UMAP represents a weighted combination of both the scRNA-seq and scATAC-seq datasets. Clustering shows pre-OPCs, OPCs, neural progenitors (NP1), pre-motor neurons (pre-MNs), and motor neurons (MN1). B) Differential gene expression analysis for pre-OPCs compared to NP1. Magenta dots represent genes enriched in pre-OPCs whereas cyan dots represent genes enriched in NP1. Genes not reaching statistical significance are indicated by gray dots. Wnt inhibitor *barhl2* is highlighted by a red box. C) Differential gene expression analysis for pre-OPCs compared to pre-MNs. Magenta dots represent genes enriched in pre-OPCs whereas cyan dots represent genes enriched in pre-MNs. Genes not reaching statistical significance are indicated by gray dots. Wnt inhibitor *cachd1* is highlighted by a red box. D) Schematic depicting the canonical Wnt signaling pathway and the distinct roles of Cachd1 and Barhl2 in Wnt inhibition. Cachd1 prevents binding of Wnt ligands to the LRP/Frizzled receptor complex whereas Barhl2 binds to TCF transcription factors and promotes a repressive state. E) Gene expression profiles of *barhl2* (left) and *cachd1* (right) on our RNA UMAP. Black boxes zoom in on the pre-OPC cluster, demonstrating expression of both genes in pre-OPCs. F) Chromatin accessibility of peaks located in DNA regions associated with *barhl2* (left) or *cachd1* (right) in pre-OPCs, pre-MNs, and neural progenitors. Blue tones indicate closed chromatin, and red tones indicate more accessible chromatin. Each column represents one chromatin peak. G) Representative images of fluorescent in situ RNA hybridization to detect *barhl2* (cyan) and *cachd1* (yellow) in transverse sections of 24, 36, and 48 hpf embryos. *olig2* (magenta) marks the pMN domain. Dotted white lines denote the spinal cord and white boxes denote the pMN domain. Cell bodies labeled by DAPI are shown in gray. Scale bars: 10 µm.

Notably, several RNA transcripts encoding proteins known to modulate Wnt signaling were enriched in pre-OPCs, including *barhl2* and *cachd1* (Figure 1B, C). Barhl2 is a transcription factor that binds and stabilizes Tcf transcription factor repressive complexes, preventing activation of downstream Wnt signaling (Bou-Rouphael et al., 2024; Bou-Rouphael & Durand, 2021; Juraver-Geslin et al., 2011; Sena et al., 2019). Cachd1 is a transmembrane protein that binds to Wnt receptors LRP/Frizzled and can interfere with Wnt ligand-receptor interactions (Ablinger et al., 2022; Powell et al., 2024; Scala et al., 2023) (Figure 1D). pre-OPCs uniquely expressed *barhl2* whereas pre-OPCs, neural progenitors, and lateral floor plate cells expressed *cachd1* (Figure 1E). To learn if this differential gene expression correlates with differences in chromatin accessibility, we utilized Signac (Stuart et al., 2021) to identify accessible chromatin regions, or peaks, in pre-OPCs, pre-MNs, and neural progenitors and assigned each peak to the gene nearest that chromatin region. We plotted the accessibility for peaks near the genes encoding *barhl2* and *cachd1* across these clusters. Chromatin peaks associated with *barhl2* or *cachd1* were more accessible in pre-OPCs compared to neural progenitors and pre-MNs (Figure 1F). From these data, we conclude that these Wnt inhibitor-encoding genes are actively remodeled to enable their expression in pre-OPCs.

To examine the expression patterns of *barhl2* and *cachd1* in vivo, we performed fluorescent in situ RNA hybridization on transverse sections of the zebrafish spinal cord to detect them simultaneously in combination with *olig2* prior to (24 hpf), during (36 hpf), and after (48 hpf) OPC specification. At 24 hpf, few spinal cord cells expressed *barhl2*. At 36 hpf, *olig2^+^* pMN cells expressed high levels of *barhl2* transcripts. By 48 hpf, spinal cord cells no longer expressed *barhl2* (Figure 1G). Similarly, spinal cord cells highly expressed *cachd1* at 36 hpf, with lower levels of *cachd1* RNA transcripts present at 24 hpf and 48 hpf (Figure 1G). Together, our multi-omics and in vivo data raise the possibility that pre-OPC expression of *barhl2* and *cachd1* is an intrinsic mechanism of Wnt inhibition that drives neural progenitors into an OPC fate.

### Wnt signaling activity declines in neural progenitors prior to OPC specification

The temporal patterns of Wnt inhibitor gene expression suggest that Wnt signaling is dynamically regulated during the period of OPC specification. To determine whether regulation of Wnt signaling by pre-OPCs might contribute to OPC formation, we first assessed endogenous Wnt signaling dynamics within the ventral spinal cord. To do so, we evaluated transverse sections of the zebrafish spinal cord at 36 and 48 hpf for active Wnt signaling. We utilized the transgene *Tg(7xTCF-Xla.Siam:nls-mcherry),* referred to in this manuscript as *Tg(7xTCF:nls-mch),* which encodes nuclear-localized mCherry fluorescent protein under the control of seven tandem repeats of TCF/Lef-binding sites fused to a minimal promoter to amplify endogenous Wnt/ß-Catenin signaling-dependent gene expression (Moro et al., 2012). We combined this transgene with *Tg(olig2:egfp)* and performed immunohistochemistry to detect Sox10 to label oligodendrocyte lineage cells. At 36 hpf, cells distributed throughout the spinal cord expressed the Wnt reporter at various levels of fluorescent intensity. Within the pMN domain, Wnt reporter expression was similarly variable, with some cells having little to no detectable expression. Particularly, very few Sox10^+^ OPCs expressed the reporter (Figure 2A). By 48 hpf, fewer cells throughout the spinal cord expressed the Wnt reporter, however the percentage of Sox10^+^ OPCs that expressed the reporter increased (Figure 2A, B). This change in Wnt activity from OPC specification to OPC differentiation is consistent with reports of a complex, stage-dependent role for Wnt signaling in OPC maturation (Dai et al., 2014; Soomro et al., 2018).

**Figure 2:**
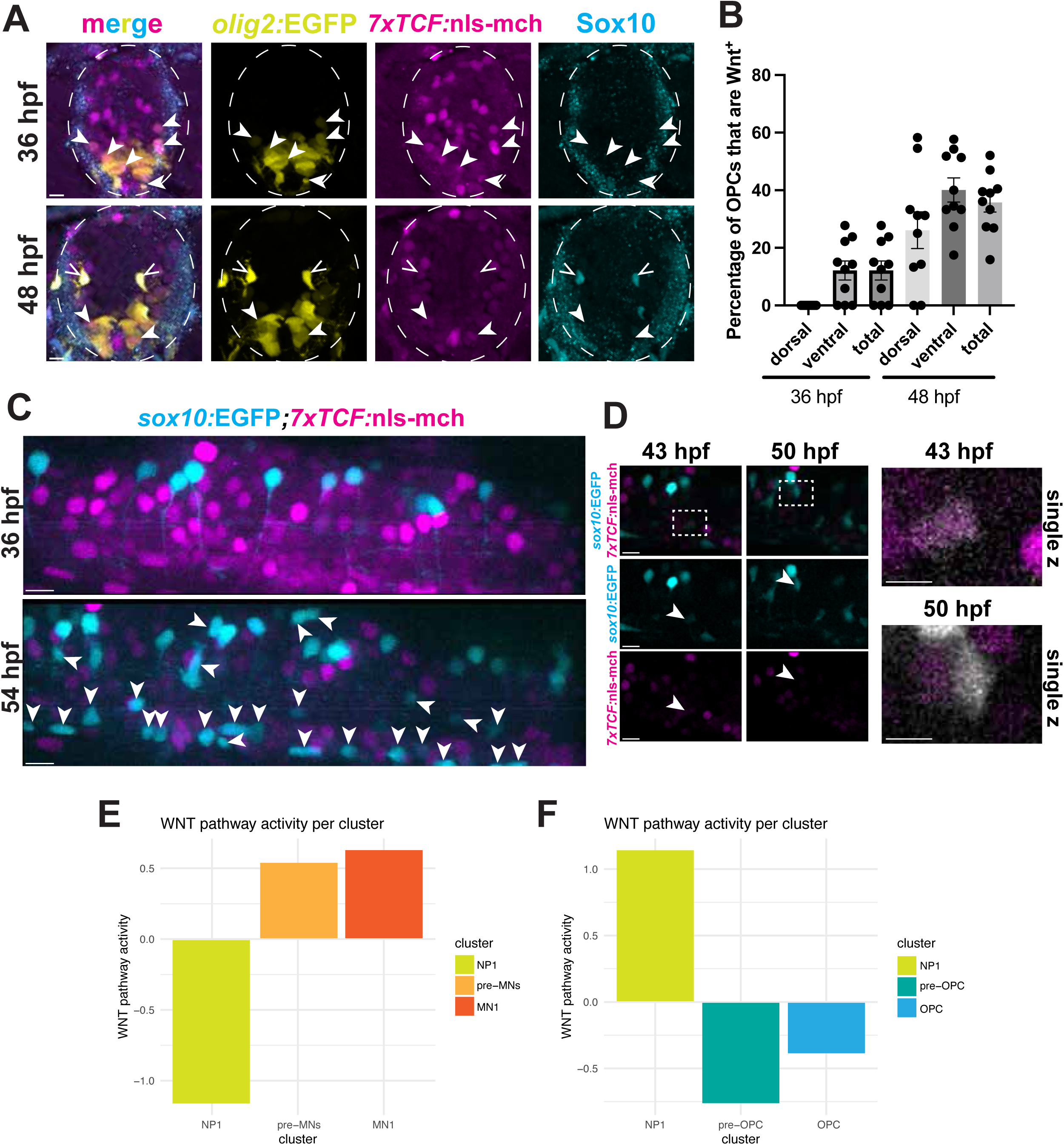
Wnt signaling activity declines in neural progenitors prior to OPC specification. A) Representative images of immunohistochemistry against Sox10 (cyan) in transverse sections of 36 hpf (top panels) and 48 hpf (bottom panels) *Tg(olig2:egfp);Tg(7xTCF:nls-mch)* embryos *olig2:*EGFP (yellow) marks the pMN domain and *7xTCF:*nls-mch (magenta) marks Wnt-active cells. Dotted white lines denote the spinal cord. Solid arrowheads indicate cells that are *olig2:*EGFP*^+^, 7xTCF:*nls-mch^+^, and Sox10^-^. Lined arrowheads indicate cells that are *olig2:*EGFP*^+^, 7xTCF:*nls-mch*^+^,* and Sox10*^+^.* Scale bars: 10 µm. B) Quantification of the percentage of *olig2^+^/*Sox10^+^ OPCs that are *7xTCF:*nls-mch*^+^* at 36 hpf (left) and 48 hpf (right). Each dot on the bar plot represents an individual embryo. C) Representative images from beginning (36 hpf) and end (54 hpf) of an embryo expressing *Tg(sox10:egfp)* (cyan) and *Tg(7xTCF:nls-mch)* (magenta). White arrows indicate newly formed OPCs at 54 hpf. Scale bars: 10 µm. D) Representative images of individual cells from (C) at the onset of *sox10* expression (43 hpf) and later after initiation of migration (50 hpf). White arrows denote the individual cells. Dotted white boxes indicate the locations of the inset boxes (right) at 43 hpf (top) and 50 hpf (bottom). Each image represents a single z-plane from the movie z-stack. *sox10:*EGFP cells are pseudo-colored gray to better demonstrate contrast with *7xTCF:*nls-mch. Scale bars: 10 µm (panels), 5 µm (insets). E) PROGENy analysis of Wnt pathway activity for the neuronal lineage (NP1, pre-MNs, MN1). F) PROGENy analysis of Wnt pathway activity for the oligodendrocyte lineage (NP1, pre-OPCs, OPC).

Whereas a subset of pMN cells had active Wnt signaling at 36 hpf, these rarely included OPCs, consistent with the possibility that Wnt is actively silenced for OPC specification. To directly observe changes in Wnt signaling as OPCs are specified, we conducted confocal light microscopy of embryos carrying the transgenes *Tg(sox10:egfp)*, in which oligodendrocyte lineage cells express EGFP under control of *sox10* regulatory DNA, and *Tg(7xTCF:nls-mch),* from 36 to 54 hpf. Consistent with our static imaging analysis, the number of *7xTCF:*nls-mch*^+^* cells declined from 36 hpf to 54 hpf (Figure 2C). Further, *7xTCF:*nls-mch expression decreased in individual cells prior to activation of *sox10:*EGFP (Figure 2D). These data provide evidence that pMN progenitors downregulate Wnt signaling prior to OPC formation.

To explore how Wnt activity might predict neural progenitor fate, we utilized our single-cell RNA sequencing and PROGENy (Schubert et al., 2018) to infer Wnt activity levels in neural progenitors, OPC populations, and motor neuron populations. PROGENy predicts Wnt pathway activity in individual cells by combining the gene expression values of Wnt-responsive genes from our single-cell RNA sequencing with pre-determined scores based on orthologous genes that indicate the influence of each gene on active pathway signaling based on prior research. This generates a score that predicts how active Wnt signaling is in a particular cluster. Wnt signaling signatures were stronger in pre-MN and motor neuron populations relative to neural progenitors (Figure 2E). Conversely, pre-OPCs and OPCs had lower Wnt activity relative to neural progenitors (Figure 2F). These data support the possibility that active inhibition of Wnt signaling promotes the specification of neural progenitors into OPCs.

### Inhibition of Wnt signaling promotes OPC specification

Our single-cell multi-omics data revealed that pre-OPCs express genes involved in Wnt inhibition (Figure 1) and in vivo analyses confirmed that Wnt signaling decreases in pMN progenitor cells prior to OPC specification (Figure 2). These data raise the possibility that Wnt inhibition promotes OPC specification. To test this, we utilized IWR-1, which facilitates degradation of ß-Catenin (Chen et al., 2009), to pharmacologically inhibit Wnt signaling. We administered 10 µM IWR-1 or 0.5% DMSO to *Tg(sox10:egfp);Tg(7xTCF:nls-mch)* embryos at 30 hpf, following the majority of motor neuron specification but prior to the onset of OPC specification. Time-lapse confocal microscopy for 22 hours starting at 36 hpf revealed reduced Wnt signaling in IWR-1-treated embryos, confirming the efficacy of the drug treatment. Further, there was a significant increase in the number of OPCs in IWR-1-treated embryos compared to DMSO-treated embryos (Figure 3A). Additionally, OPCs formed slightly earlier in IWR-1-treated embryos compared to DMSO-treated embryos (Figure 3B). The increase in OPC numbers was significant in both the dorsal and ventral spinal cord at 58 hpf (Figure 3C). To determine whether this increase in OPCs was due to enhanced proliferation of newly formed OPCs, we quantified the number of cell divisions that occurred in Sox10^+^ cells during our time-lapse movies. There was no significant change in OPC cell divisions (Figure 3D). Therefore, pharmacological inhibition of Wnt leads to an overall increase in the number of OPCs produced by neural progenitors.

**Figure 3:**
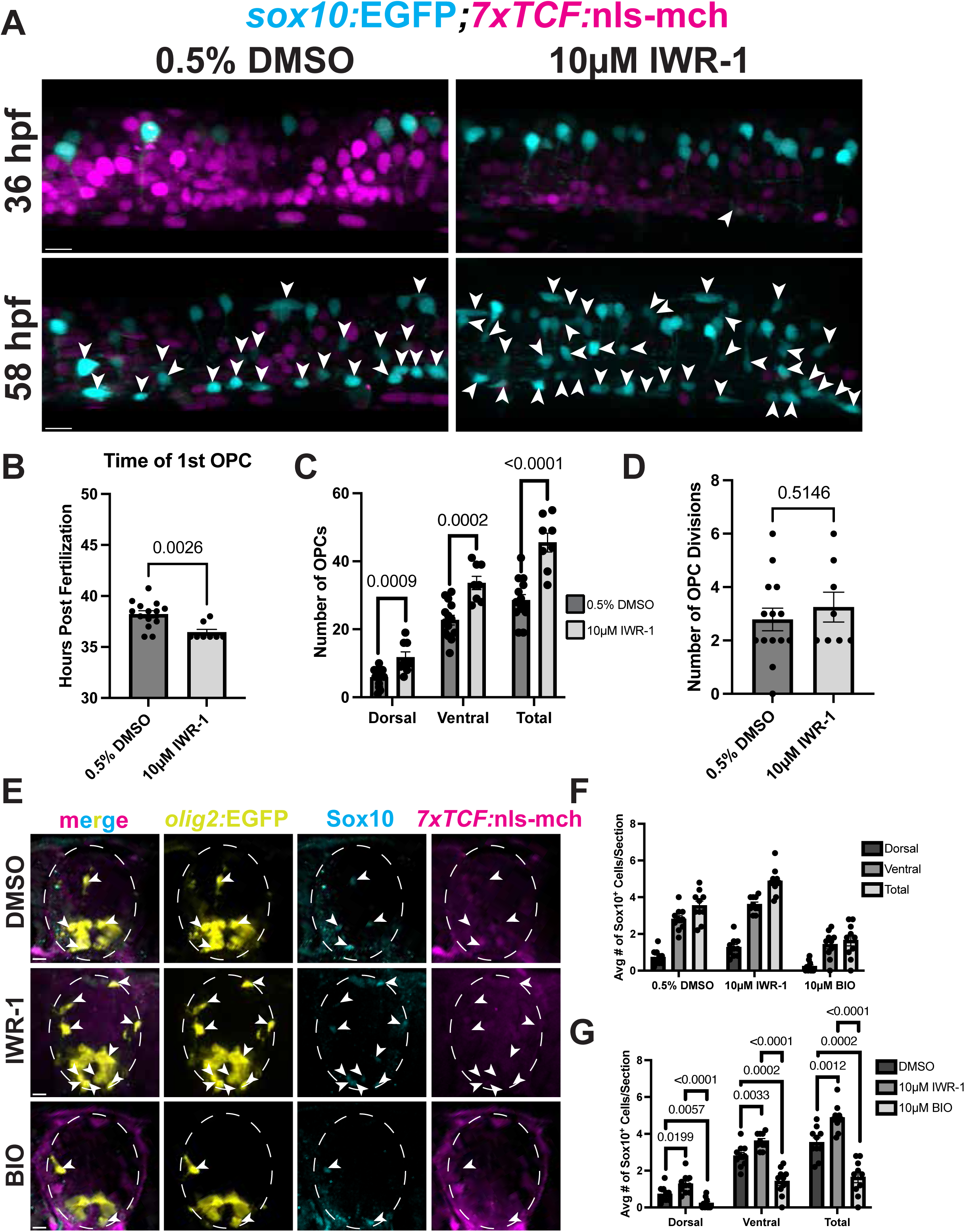
Inhibition of Wnt signaling promotes OPC specification. A) Representative images from the starting frame (36 hpf) and ending frame (58 hpf) of 22-hour time-lapse movies of embryos expressing *sox10:*EGFP (cyan) and *7xTCF:*nls-mch (magenta). Embryos were treated with either 0.5% DMSO (left panels) or 10 µM IWR-1 (right panels) at 30 hpf and throughout imaging. White arrowheads depict *sox10^+^* OPCs. Scale bars: 10 µm. B) Quantification of the time of the first *sox10^+^* OPC appearance in 0.5% DMSO or 10 µM IWR-1 treated embryos. C) Quantification of the number of *sox10^+^* OPCs present in the dorsal and ventral spinal cord at 58 hpf. D) Quantification of the number of cell divisions for *sox10^+^* OPCs. For (B-D), each dot represents an individual embryo. Unpaired t-tests were performed to determine statistical significance between groups. E) Representative images of immunohistochemistry against Sox10 (cyan) in transverse sections of 48 hpf *Tg(olig2:egfp);Tg(7xTCF:nls-mch)* embryos treated with either 0.5% DMSO (top panels), 10µM IWR-1 (middle panels), or 10µM BIO (bottom panels). White arrowheads depict Sox10/*olig2*^+^ OPCs. Scale bars: 10µm. F) Quantification of the average number of Sox10/*olig2^+^*OPCs per section in the dorsal and ventral spinal cord for 0.5% DMSO, 10µM IWR-1, and 10µM BIO-treated groups, with each dot representing an individual embryo. G) Quantifications from E) and F) with statistical analyses comparing the average number of Sox10/*olig2*^+^ cells per section in the dorsal and ventral spinal cord for each treated group. Unpaired t-tests were performed between each treatment group at each spinal cord location to determine statistical significance.

To validate our live-imaging findings, we obtained transverse sections of 48 hpf 10 µM IWR-1 or 0.5% DMSO-treated *Tg(olig2:egfp);Tg(7xTCF:nls-mch)* embryos and conducted immunohistochemistry to detect Sox10. Consistently, IWR-1-treated embryos produced excess OPCs occupying both the dorsal and ventral spinal cord compared to DMSO-treated embryos (Figure 3E-G), demonstrating that excess OPCs are specified in the absence of Wnt signaling. Because our pharmacological inhibition of Wnt signaling increased OPC production, we predicted that elevated Wnt activity would suppress OPC formation. To test this, we treated 30 hpf *Tg(olig2:egfp);Tg(7xTCF:nls-mch)* embryos with 10 µM BIO, a Wnt agonist that inhibits GSK-3 and stabilizes ß-catenin (Meijer et al., 2003), to induce Wnt signaling during OPC specification. We produced transverse sections of 48 hpf 10 µM BIO or 0.5% DMSO-treated embryos and conducted immunohistochemistry to detect Sox10. BIO-treated embryos had significantly fewer OPCs compared to DMSO-treated embryos (Figure 3E-G). Together, these data reveal that modulation of Wnt signaling regulates the timing and number of OPCs specified during spinal cord development.

To investigate whether the Wnt inhibitors expressed by pre-OPCs are necessary for OPC specification (Figure 1), we used CRISPR/Cas9 mutagenesis to establish *barhl2* (Figure 4A) and *cachd1* (Figure 4B) loss-of-function alleles. For embryos homozygous for mutations of either gene, there were no obvious changes in larval morphology or lethality. To assess OPC numbers, we produced transverse sections of *barhl2^-/-^* and *cachd1^-/-^*embryos and their wild-type siblings carrying the *Tg(olig2:egfp)* transgene at 48 hpf and performed immunohistochemistry to detect Sox10. Both *barhl2^-/-^* and *cachd1^-/-^* embryos produced significantly fewer OPCs than their wild-type siblings (Figure 4C-F). These data demonstrate that Wnt inhibitors uniquely expressed by pre-OPCs act as positive regulators of OPC specification.

**Figure 4:**
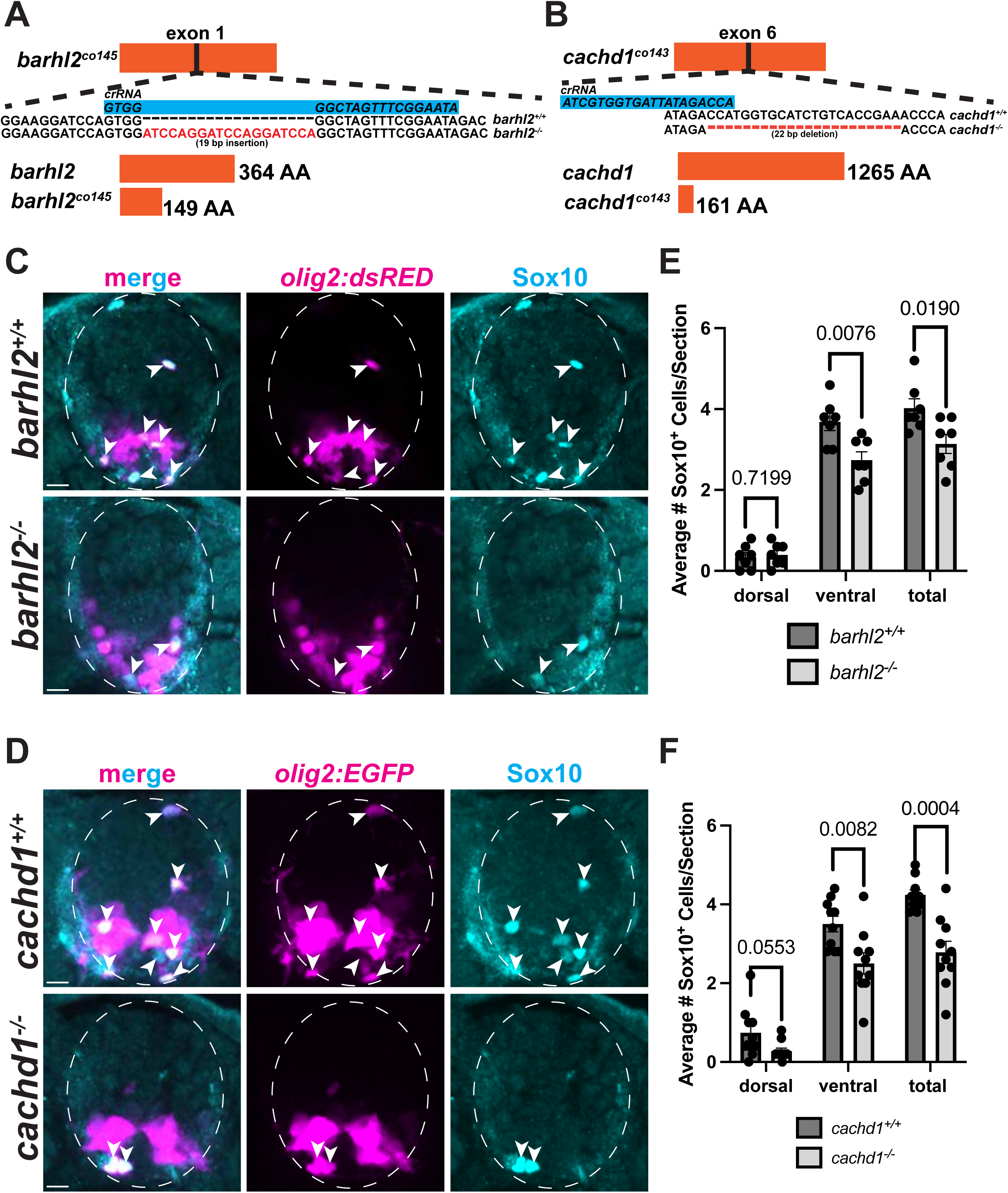
Deletion of pre-OPC-specific Wnt inhibitors decreases OPC production. A) Schematic describing the 19 bp insertion in the *barhl2^co145^* allele which results in an early stop codon and disrupted protein function. B) Schematic describing the 22 bp deletion *cachd1^co143^* allele which results in an early stop codon and disrupted protein function. Representative images of immunohistochemistry against Sox10 (cyan) in transverse sections of 48 hpf *barhl2^+/+^* and *barhl2^-/-^* embryos (C) or *cachd1^+/+^* and *cachd1^-/-^* embryos carrying the *Tg(olig2:egfp)* transgene (D). Arrowheads depict Sox10/*olig2*^+^ OPCs. Scale bars: 10 µm. E-F) Quantification of the average number of dorsal and ventral Sox10/*olig2*^+^ OPCs per section from *barhl2* (E) or *cachd1* (F) experiments (C,D), with each dot representing an individual embryo. Unpaired t-tests were performed to determine statistical significance between wild-type and mutant embryos.

### Ventral spinal cord cells express *wnt4b,* which regulates OPC formation

To identify the sources and identities of Wnt ligands important for OPC formation, we utilized our single-cell RNA sequencing (Arena et al., 2026) to explore which Wnt ligands are expressed by ventral spinal cord cells, close to the origin of OPCs. Cells identified as lateral floor plate expressed *wnt4b*, consistent with prior literature (Inohaya et al., 2010; Liu et al., 2000),whereas lateral floorplate cells and OPCs expressed *wnt7aa* (Figure 5A-C). To confirm the expression of these ligands in vivo, we conducted fluorescent RNA in situ hybridization on transverse sections of the zebrafish spinal cord to detect both *wnt4b* and *wnt7aa* at 24, 36, and 48 hpf (Figure 5D). Ventral spinal cord cells in the position of the lateral floor plate expressed *wnt4b* from 24 through 48 hpf and *wnt7aa* at 48 hpf (Figure 5D). The time and location of *wnt4b* expression is consistent with the possibility that it regulates OPC specification.

**Figure 5:**
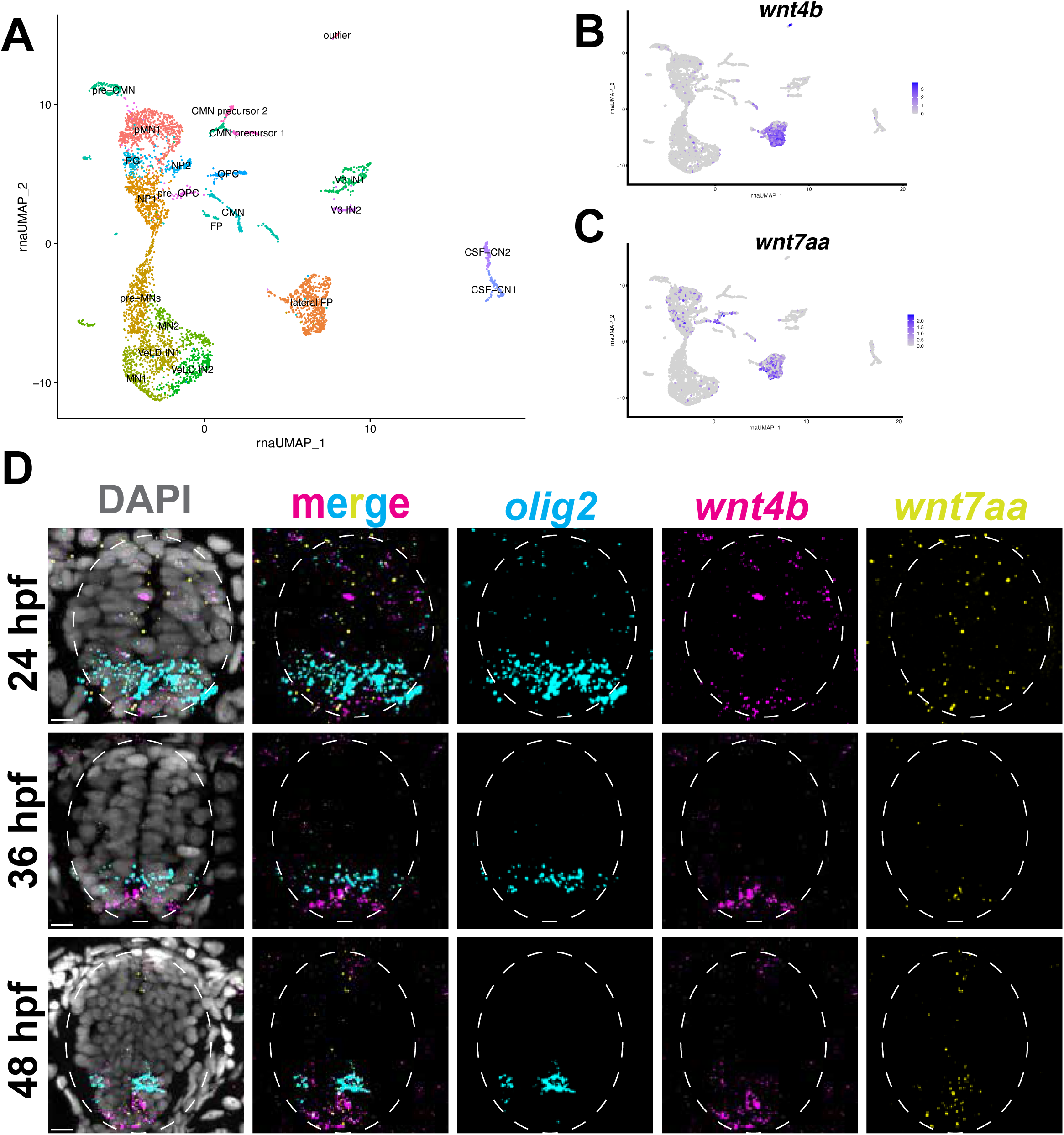
Ventral spinal cord cells express *wnt4b* and *wnt7aa*. A) RNA UMAP generated from our scRNA-seq multi-omics dataset. B) RNA UMAP for *wnt4b*. C) RNA UMAP for *wnt7aa*. D) Representative images of fluorescent in situ RNA hybridization to detect *olig2* (cyan), *wnt4b* (magenta), and *wnt7aa* (yellow) in transverse sections of 24, 36, and 48 hpf embryos. *olig2* (cyan) marks the pMN domain. Dotted white lines denote the spinal cord. Cell bodies labeled by DAPI are shown in gray. Scale bars: 10 µm.

To investigate if *wnt4b* provides the Wnt activity that suppresses OPC formation, we utilized CRISPR/Cas9 gene editing to create loss of function *wnt4b* F0 embryos, referred to as “crispants”. To assess OPC numbers, we produced transverse sections of 48 hpf control or *wnt4b* crispant embryos expressing *Tg(olig2:egfp)* and performed immunohistochemistry to detect Sox10. *wnt4b* crispant embryos had significantly more OPCs compared to control siblings (Figure 6). These data demonstrate that in the absence of *wnt4b*, OPC specification is enhanced, suggesting *wnt4b* as a source of ventral Wnt that must be inhibited for OPC specification to occur.

**Figure 6:**
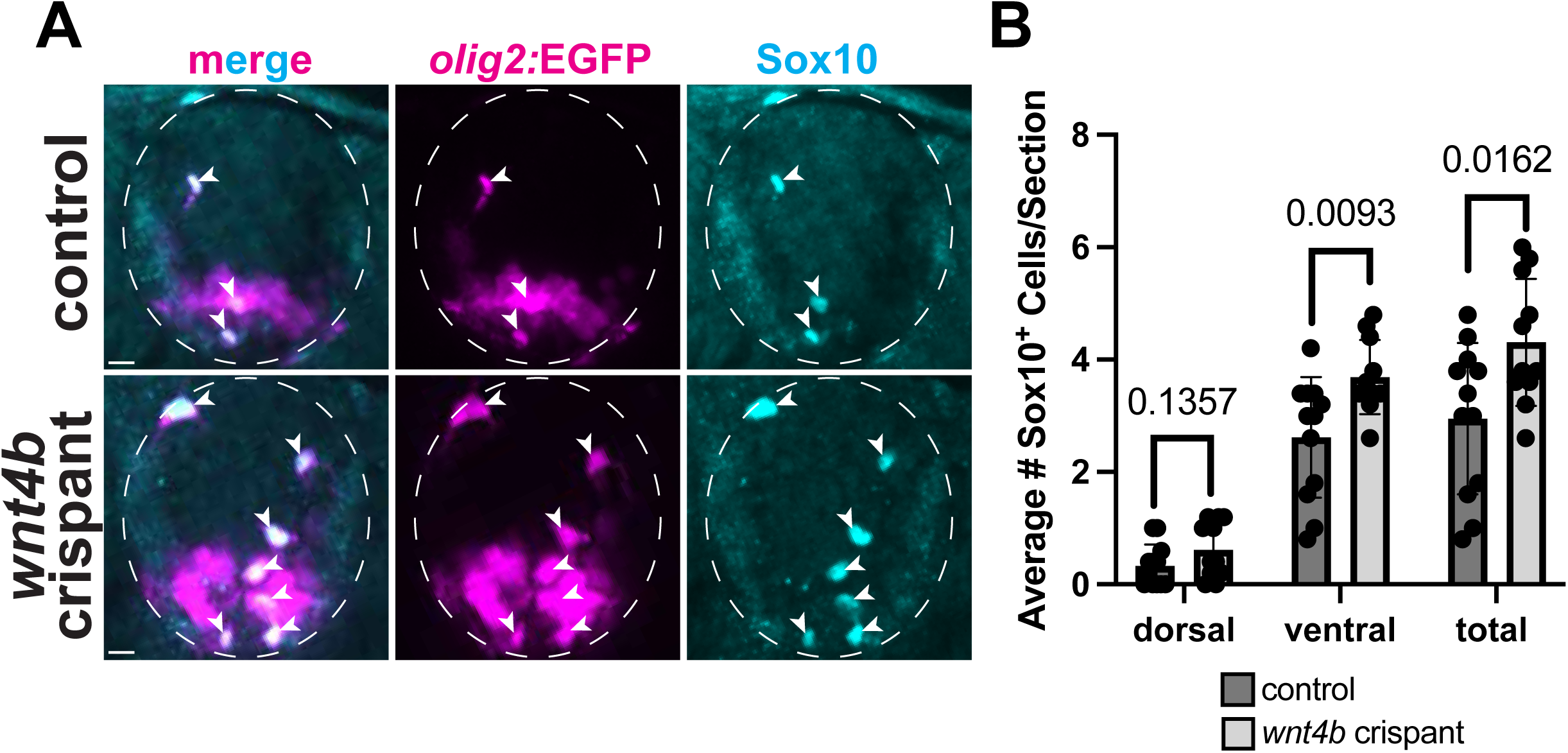
Disruption of *wnt4b* enhances OPC production. A) Representative images of immunohistochemistry against Sox10 (cyan) in transverse sections of 48 hpf *Tg(olig2:egfp)* control (top panels) or *wnt4b* crispant (bottom panels) embryos. Arrowheads depict Sox10/*olig2*^+^ OPCs. Dotted white lines denote the spinal cord. Scale bars: 10 µm. B) Quantification of the average number of dorsal and ventral Sox10/*olig2*^+^ OPCs per section in control or *wnt4b* crispant embryos, with each dot representing an individual embryo. Unpaired t-tests were performed to determine statistical significance between control and *wnt4b* crispant embryos.

### A gene regulatory network for intrinsic inhibition of Wnt signaling in pre-OPCs

Our data are consistent with a model whereby pre-OPC inhibition of ventrally derived Wnt4b function drives a subset of pMN progenitors into the oligodendrocyte lineage. We identified *barhl2* and *cachd1* as genes encoding Wnt inhibitors that are expressed by pre-OPCs, however it remains unclear how transcription of these genes is activated. To explore mechanisms that potentially regulate transcription of *barhl2* and *cachd1*, we utilized our single-cell multi-omics dataset to identify regions of open chromatin associated with these gene loci. For both genes, we utilized Seurat (Hao et al., 2023) and Signac (Stuart et al., 2021) to identify open chromatin regions, “peaks”, that were specific to the pre-OPC cluster (Figure 8A). Further, we computed statistical correlations between the accessibility of individual open chromatin peaks and the RNA transcript expression of the gene in our pre-OPC cluster and identified “linked peaks” that are likely to be associated with gene expression (Figure 7A). We identified the correlating sequence for these linked peaks and confirmed that their accessibility is specific to pre-OPCs (Figure 7B, C). Next, we scanned these linked peaks for statistically enriched transcription factor binding motifs using JASPAR (Ovek Baydar et al., 2025). This analysis identified potential transcription factors that could bind at *barhl2* or *cachd1* linked peaks. We then plotted the top ten transcription factors with enriched motifs at linked peaks for *barhl2* (Figure 7D) and *cachd1* (Figure 7E). Notably, Gsx2, which we previously identified as important for OPC specification (Arena et al., 2026), had an enriched motif at the *barhl2* linked peak. Further, Gli3, an active component of the Shh pathway that can negatively regulate Wnt signaling (Tyurina et al., 2005; Ulloa et al., 2007), had an enriched motif at the *cachd1* linked peak. Together, these analyses provide evidence for transcriptional activation of Wnt inhibitors in pre-OPCs.

**Figure 7:**
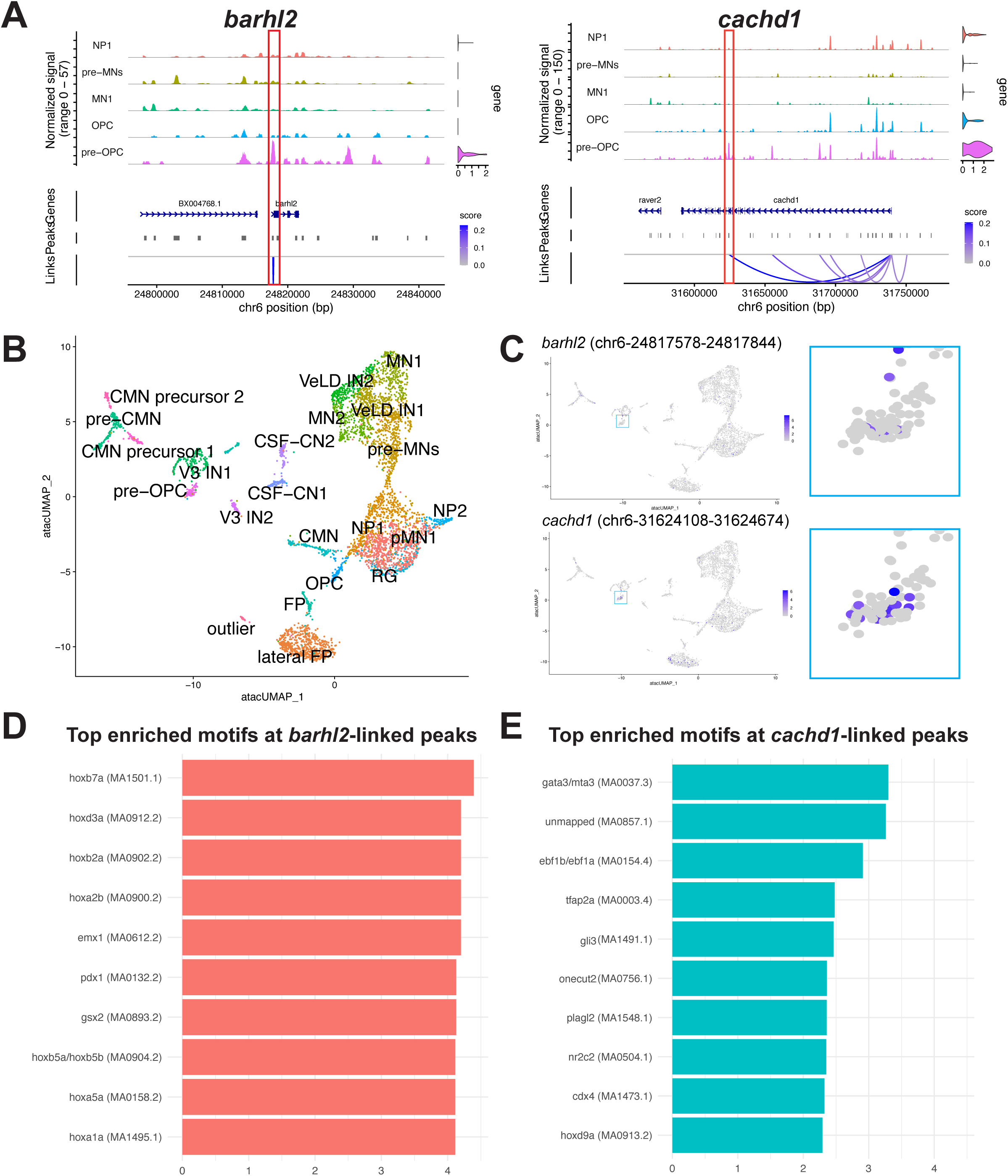
Single-cell multi-omic analyses predicting a gene regulatory network for pre-OPC expression of *barhl2* and *cachd1*. A) Linked peak plots for *barhl2* (left) and *cachd1* (right) at 36 hpf for NP1 (orange), pre-MNs (yellow), MN1 (green), OPC (blue) and pre-OPC (magenta) clusters. Violin plots of *barhl2* and *cachd1* expression are on the right. Peaks (middle) indicate regions of chromatin accessibility and map to genomic locations (bottom). Individual peaks linked with gene expression are indicated by blue lines, with darker blue representing a stronger correlation between peak accessibility and gene expression. Red boxes indicate strongly linked peaks for both genes. B) ATAC UMAP generated from our scATAC-seq mutli-omics dataset. C) ATAC UMAP for the highlighted linked peaks in A) for *barhl2* (top) and *cachd1* (bottom). Insets (turquoise box) highlight peak accessibility in the pre-OPC cluster. D-E) Top enriched motifs for transcription factors (JASPAR) that can bind at the highlighted *barhl2* (D) and *cachd1* (E) linked peaks.

## DISCUSSION

In the developing spinal cord, morphogen gradients establish combinatorial transcription factor codes that pattern the dorsal-ventral axis. Wnt/BMP signaling originating from the roof plate opposes notochord-derived Shh signaling, creating progenitor subdomains that give rise to specific subsets of neurons and glia (Brafman & Willert, 2017; Freese et al., 2009; Guo et al., 2015; Megason & McMahon, 2002; Soomro et al., 2018). In the pMN domain of the ventral spinal cord, NPCs produce neurons followed by oligodendrocytes. Prior research has characterized Wnt signaling as a negative regulator of oligodendrogenesis (Dai et al., 2014; Langseth et al., 2010) and demonstrated that Wnt activity declines as neurogenesis shifts to gliogenesis (Guo et al., 2015; Langseth et al., 2010; Soomro et al., 2018). However, these studies did not address the mechanisms through which Wnt signaling is dampened to drive oligodendrocyte specification. Here, we reveal that a transcriptionally distinct subpopulation of NPCs in the zebrafish spinal cord, pre-OPCs, express genes encoding Wnt inhibitors Barhl2 and Cachd1. We show that inhibition of Wnt signaling enhances OPC production in vivo and that *barhl2* and *cachd1* positively regulate OPC specification, supporting a model where cell-intrinsic Wnt inhibition specifies pre-OPCs for oligodendrocyte fate.

Zebrafish pre-OPCs express both *barhl2* and *cachd1*, genes encoding two distinct Wnt inhibitors. Cachd1 is a highly conserved transmembrane protein that simultaneously binds Lrp6 and Frizzled family Wnt co-receptors. By bridging these receptors, Cachd1 physically competes with endogenous Wnt ligands (Powell et al., 2024; Scala et al., 2023). This receptor-level interference prevents downstream intracellular stabilization of ß-catenin and expression of target genes. Therefore, *cachd1* expression by pre-OPCs desensitizes them to surrounding Wnt signals. In parallel, homeobox transcription factor Barhl2 suppresses Wnt activation by stabilizing the Tcf7l1-Groucho transcriptional co-repressor complex and recruiting Hdac1 to chromatin to repress target genes (Sena et al., 2019). Expression of both *barhl2* and *cachd1* highlights dual mechanisms of intrinsic Wnt inhibition that target distinct components of the Wnt signaling pathway. Production of oligodendrocytes following neurogenesis is essential for proper myelination of developing axons and for establishing efficient neural circuits. Therefore, it would reasonably be advantageous for organisms to develop redundant mechanisms safeguarding the specification of oligodendrocytes. Accordingly, we found that loss of either *barhl2* or *cachd1* reduced, but did not eliminate, OPC production. However, neither *barhl2* nor *cachd1* mutants reduced OPC specification to the degree that treatment with Wnt agonist BIO did, suggesting that redundant in vivo mechanisms of Wnt inhibition ensure OPC generation.

Canonical Wnt signaling is widely recognized as one of the primary morphogens responsible for patterning the dorsal-ventral axis of the developing vertebrate spinal cord (Brafman & Willert, 2017; Megason & McMahon, 2002; Muroyama et al., 2002). Wnt ligands are typically distributed in a high-dorsal to low-ventral gradient (Brafman & Willert, 2017; Megason & McMahon, 2002), and OPC specification is mostly restricted to the ventral spinal cord (Warf et al., 1991). Oligodendrogenesis coincides with a decline in the Wnt gradient, suggesting lower availability of Wnt ligands drives cells toward a glial fate (Guo et al., 2015). However, prior research has documented floor plate expression of Wnt ligands at various stages of early development in zebrafish, medaka, chick, and mouse models (Agalliu et al., 2009; Inohaya et al., 2010; Liu et al., 2000), suggesting ventral Wnt sources could contribute to pMN cell fate decisions. Accordingly, we observed that RNAs encoding Wnt ligands Wnt4b and Wnt7aa are highly abundant ventrally during oligodendrogenesis. Further, we determined that overall Wnt activity declines across the spinal cord from 24 hpf to 48 hpf despite persistence of *wnt4b* and *wnt7aa* expression. This indicates that the correlation between low Wnt activity and glial fate commitment is not simply a product of reduction in extracellular ligand availability, but rather a transition in which NPCs actively become unresponsive to Wnt. This finding aligns with previous observations of NPCs losing their ability to respond to Wnt signaling around E9.5 in mice, prior to oligodendrogenesis at E12.5 (Yu et al., 2008). Finally, we found that loss of functional *wnt4b* leads to an increase in OPCs, confirming that this ligand can regulate OPC generation. Together, these data support a model where ventral Wnt ligands act as local repressors of gliogenesis, with pre-OPCs expressing Wnt inhibitor genes to permit oligodendrocyte specification.

Our pharmacological manipulations of Wnt signaling during OPC specification corroborate a model where Wnt inhibition promotes oligodendrogenesis. These findings complement a recent study in mice that shows Wnt activation suppresses early oligodendrocyte generation by directly downregulating Olig2 (Jiang et al., 2020). However, several studies have characterized a positive role for Wnt signaling in oligodendrocyte maturation and myelination later in development (Dai et al., 2014; Fancy et al., 2009; Guo et al., 2015; Soomro et al., 2018; Tawk et al., 2011). Accordingly, we observed a decrease in *barhl2* and *cachd1* in vivo at 48 hpf, following OPC specification. Similarly, our single-cell RNA sequencing shows high expression of *barhl2* and *cachd1* in pre-OPCs with expression levels declining in OPCs. However, expression of *tcf7l2*, a Wnt-responsive transcription factor previously implicated in oligodendrocyte development (Fu et al., 2012; Guo & Wang, 2023; H. Wang & Matise, 2016; Zhang et al., 2021; Zhao et al., 2016), is present in both pre-OPCs and OPCs. Further, *wnt4b* and *wnt7aa* remain present in the ventral spinal cord at 48 hpf. Together, these data suggest that cell-intrinsic mechanisms for Wnt modulation are essential for OPC specification, however, the mechanisms that modulate Wnt signaling in later stages of oligodendrocyte development remain incompletely understood.

Pre-OPCs represent a transcriptionally distinct subset of NPCs that appear fated for the oligodendrocyte lineage. Our single-cell multi-omics revealed that *barhl2* and *cachd1* are differentially expressed by pre-OPCs and the chromatin regions associated with these genes have unique accessibility peaks. To understand how pre-OPC-specific expression of Wnt inhibitors is initiated, we utilized our single-cell multi-omics dataset to explore potential transcriptional regulators of *barhl2* and *cachd1*. We identified *gsx2* as one likely regulator of *barhl2.* We recently characterized *gsx2* as uniquely and transiently expressed by pre-OPCs, where it serves as a critical temporal regulator of OPC formation (Arena et al., 2026). *gsx2* is highly expressed by pre-OPCs and is downregulated as specification progresses, making its gene expression profile similar to that of *barhl2*. However, loss of *gsx2* leads to excess OPCs, suggesting *gsx2* holds pre-OPCs in a progenitor state until they are ready to progress into OPCs. Conversely, loss of *barhl2* leads to a reduction in OPCs. This suggests that *gsx2* might repress expression of *barhl2* until it is time for specification to begin. Additionally, *gli3* was identified as a probable regulator of *cachd1.* Gli3 is a critical bi-functional regulator of Shh signaling (Tyurina et al., 2005). Gli3 regulation of *cachd1* could link Shh signaling, a pathway essential for oligodendrocyte development (Nery et al., 2001; S. Oh et al., 2005; Tekki-Kessaris et al., 2001; L. C. Wang & Almazan, 2016), to cell-intrinsic Wnt inhibition. Shh signaling suppresses nuclear Gli3 repressor activity (Gli3R), allowing active Gli3 to target gene activation (S. O. Oh et al., 2009; Tyurina et al., 2005; Ulloa et al., 2007). This raises the possibility that Shh-dependent modulation of Gli3 could provide a mechanism for activation of *cachd1* in pre-OPCs. Together, these data highlight the complex regulatory mechanisms that dictate cell-responsiveness to Wnt signaling to drive cell fate decisions in the developing ventral spinal cord.

Our study has some important limitations. First, we cannot track the fates of neural progenitors in our experiments that change the level of Wnt signaling. In zebrafish, pre-OPCs represent a small, transient subset of pMN progenitors, and we have not yet created pre-OPC-specific transgenic reporters for cell fate analyses. Because there are few pre-OPCs and OPCs relative to the number of neurons that arise from pMN progenitors, any change in neuron number that occurs concomitantly with changes in OPC number is difficult to detect. Consequently, we do not know if interfering with Wnt inhibition produces excess OPCs because some progenitors are diverted from neuronal fates. A second limitation is that, although we identified *wnt4b* as a ventral source of Wnt signaling for regulating OPC production, we do not know that it is the only Wnt acting on pMN progenitors. However, the effect on OPC number of *wnt4b* CRISPR/Cas9 knockdown is nearly the same as the effect of IWR-1 treatment, suggesting that Wnt4b is the primary Wnt acting to regulate OPC formation in the zebrafish spinal cord.

## MATERIALS AND METHODS

### Zebrafish Lines and Husbandry

All animal work was approved by the Institutional Animal Care and Use Committee (IAUCUC) at the University of Colorado Anschutz. Zebrafish strains utilized in this study include: AB (ZIRC), *Tg(olig2:egfp)^vu12^* (Shin et al., 2003), *Tg(olig2:dsRed2)^vu19^*(Kucenas et al., 2008), *Tg(sox10:egfp)^ba2^* (Carney et al., 2006), and *Tg(7xTCF-Xla.Siam:nls-mcherry)^ia5^* (Moro et al., 2012). Embryos were raised in petri dishes at 28.5°C in egg water (6g Instant Ocean in 20L miliQ water) and staged according to hours post fertilization (hpf) and morphological features (Kimmel et al., 1995). Sex cannot be determined at zebrafish embryonic and larval stages. For embryo injections, eggs were collected within 30 min of breeding and injected at the single-cell stage. Embryos fixed for experiments were euthanized with 4% Tricaine mesylate prior to whole fixation in 4 % paraformaldehyde in 1X PBS (PFA). When embryos required genotyping, tail clips were taken from fixed embryos and lysed.

### CRISPR/Cas9 Mutagenesis

We utilized the CRISPR-Cas9 system to generate our *cachd1* and *barhl2* loss-of-function mutants. We used Integrated DNA Technologies (IDT) to select the top predesigned Alt-R CRISPR-Cas9 guide RNAs (gRNAs) that target the coding sequence of exon 6 for *cachd1* and exon 1 for *barhl2* (see key resources table for gRNA sequences). To prepare our gRNAs and synthesize the ribonucleoprotein (RNP) complex, we followed the “Alt-R CRISPR-Cas9 System: *in vitro* cleavage of target DNA with ribonucleoprotein complex” protocol (https://sfvideo.blob.core.windows.net/sitefinity/docs/default-source/protocol/alt-r-crispr-cas9-protocol-in-vitro-cleavage-of-target-dna-with-rnp-complex.pdf?sfvrsn=88c43107_2, 2017). The RNP complex was created immediately prior to injections by combining 0.5 μg Cas9 (IDT) diluted in Cas9 working buffer (20 mM HEPES, 150 mM KCl, pH 7.5) with 100 ng of each gRNA and incubating the complex at 37°C for 10 min. The RNP complex was kept at room temperature until injected, where 2–3 nL was injected into wild-type embryos at the single-cell stage. F0 injected embryos were screened for insertions and/or deletions created by DNA cutting and repair with PCR primers designed specifically for the gRNA target site (see key resources table for PCR primer sequences). F0 larvae were raised to adulthood and individually outcrossed to wild-type fish. F1 embryos were anesthetized and genomic DNA was extracted using 50 mM NaOH followed by 1M Tris, pH 9.0 neutralization. Several individual F1 animals from at least two unique F0 founders were screened for mutant alleles using the same PCR primers described above. Mutant alleles were sequenced (Quintara Biosciences) and aligned with the published genome using SnapGene to identify function-breaking mutations. F1 adults carrying the selected *cachd1^co143^* or *barhl2^co145^* mutant allele were outcrossed to AB, *Tg(olig2:egfp)^vu12^,* or *Tg(olig2:dsred)^vu19^* fish and the F2 larvae were raised for experiments. All wild-type and mutant embryos were genotyped and alleles were confirmed prior to their use in experiments. *cachd1^co143^* is a 22 bp deletion in exon 6 that results in a premature stop codon, truncating the protein from 1265 amino acids to 161 amino acids. *barhl2^co145^*is a 19 bp insertion in exon 1 that results in a premature stop codon, truncating the protein from 364 amino acids to 137 amino acids.

For *wnt4b* mutagenesis, F0 injected embryos, crispants, were utilized for experiments and analysis. From a single clutch of embryos, half were injected with two *wnt4b* predesigned Alt-R CRISPR-Cas9 gRNAs (IDT) + Cas9 and half were injected with Cas9 alone as a control (see key resources table for gRNA sequences). All crispant embryos were tail clipped following data collection and insertions and deletions created by DNA cutting and repair were confirmed for each gRNA with PCR primers designed specifically for the gRNA target sites (see Key Resources table for PCR primer sequences). Our crispant embryos showed ∼85% efficiency of DNA cutting and repair resulting in insertions or deletions. All crispant embryos selected for experimentation contained insertions or deletions induced by both gRNAs.

### Immunohistochemistry

Embryos were fixed at indicated time points in 4% PFA, rocking at room temperature for 3 hours. Embryos were rinsed in 0.1% Triton/1x PBS (PBSTx) and stored at 4°C. Embryos were embedded in 1.5% agar/30% sucrose overnight at 4°C. Blocks were frozen on dry ice and 15 μm transverse sections were taken with a cryostat microtome and collected on polarized slides. Slides were mounted in Sequenza racks (Ted Pella cat 36105), washed 3 × 5 min with PBSTx, and blocked for 1 hour in 2% goat serum/2% bovine serum albumin/PBSTx before being placed in primary antibody (mixed in block) overnight at 4°C. Primary antibodies used in this paper include: rabbit anti-Sox10 (1:500 (Park et al., 2005),) and mouse anti-Isl1 (1:100, DSHB). Sections were washed for 1.5 hours in PBSTx, then incubated in secondary antibody (1:250, in antibody block) for 2 hours at room temperature. Secondary antibodies used in this paper include: AlexaFluor 568 goat anti-rabbit and AlexaFluor 568 goat anti-mouse (cLife Technologies). Sections were washed for 1 hour continuously in PBSTx and then mounted in Vectashield (Vector Laboratories, H-1000-10). For imaging analysis, fixed zebrafish sections were acquired using a 20x objective plus 1.6x optovar on a Zeiss Cell Observer SD 25 spinning disk confocal system (Carl Zeiss). Z stacks images were captured on Zen software (Carl Zeiss) and were processed and analyzed using Fiji/ImageJ. 10 sequential images were captured for each embryo and averaged per embryo for statistical analyses.

### Fluorescent in situ RNA Hybridization

Fluorescent in situ RNA hybridization (FISH) was performed using RNAscope Multiplex Fluorescent V2 Detection Kit (ACD, 323110). Zebrafish embryos were fixed at either 2 dpf or 3 dpf with 4% PFA, embedded as described above, and 15 μm cryosections were processed using manufacturer’s protocol and synthesized probes for *olig1* Ch1, *cachd1* Ch3, *barhl2* Ch2, *wnt4b* Ch2, or *wnt7aa* Ch3, followed by Opal dyes 520 (Akoya, FP1287001), 570 (Akoya, FP1488001) or 650 (Akoya, FP1496001). For imaging analysis, Z stack images were acquired from fixed zebrafish sections using Zen software with 20x objective plus 1.6x optovar on a Zeiss Cell Observer SD 25 spinning disk confocal system (Carl Zeiss). Images were analyzed in Imaris Viewer software (version 10.2).

### Pharmacological Treatments

Pharmacological treatments were performed in 24-well plates with up to five embryos per well. 30 hpf embryos were incubated in either 1.5 mL of 10 µM IWR-1 (Sigma-Aldrich, 681669), a selective inhibitor of Wnt, in 0.5% DMSO in egg water, 1.5 mL of 10 µM BIO (Sigma-Aldrich, 1686), a selective Wnt agonist, in 0.5% DMSO in egg water, or 0.5% DMSO in egg water at 28.5 °C in dark conditions. Embryos used for cryo-sectioning were treated for 18 hours and fixed in 4% PFA at 48 hpf. Embryos used for live imaging were treated for 6 hours prior to imaging and were given fresh pharmacological solutions prior to being mounted for live imaging. Tricaine was added to fresh IWR-1, BIO, or DMSO solutions for live imaging and embryos were submerged in those solutions for the entirety of imaging.

### In vivo Imaging

All embryos used for imaging were staged (Kimmel et al., 1995) and screened for transgene fluorescence. At specified stages, embryos were anesthetized using 3-aminobenzoic acid ester (Tricaine) and mounted in 0.8% low-melting point agarose in four-well glass bottom 35mm Petri dishes (Fisher, Greiner Bio-One). One to four embryos were mounted per well. Following mounting, Petri dishes were filled with egg water (6g Instant Ocean in 20L miliQ water) containing Tricaine. For embryos that were pharmacologically treated, Petri dishes were filled with egg water containing their specified pharmacological treatment and Tricaine. In vivo imaging was conducted on a Zeiss Cell Observer SD 25 spinning disk confocal system with a 40x water-immersion objective (Carl Zeiss). Time-lapse movies were set to image in 15-minute intervals for 20-22 hours. Z-stacks of ∼40 µm were collected for each image in a movie. The same approximate region of the spinal cord was imaged for each embryo. Time-lapse movies were processed and analyzed in Imaris Viewer software (version 10.2) and Fiji/ImageJ.

### Quantifications and Statistical Analyses

#### Image Blinding

The genotypes or pharmacological treatment conditions of fixed slides were blinded prior to imaging and were unblinded once analysis was complete. For live-imaging, files were blinded to pharmacological treatment conditions after imaging was complete but prior to any analysis. Files were unblinded once analysis was complete.

#### Cell Counts (Immunohistochemistry)

Cells were counted in transverse sections of *Tg(olig2:egfp)* embryos processed for immunohistochemistry as described above to detect Sox10. The number of dorsal and ventral Sox10^+^ cells from 10 sequential sections were counted per embryo and a minimum of 10 individual embryos were utilized per experimental group. Only Sox10^+^ cells that were also *olig2^+^* were counted. The number of Sox10^+^ cells per embryo were determined by averaging dorsal, ventral, and total numbers across 10 sequential sections.

#### Cell Counts (Time-lapse Movies)

To determine the number of OPCs produced, *sox10^+^* cells were counted in time-lapse movies of *Tg(sox10:egfp);Tg(7xTCF-Xla.Siam:nls-mcherry)* embryos at 58 hpf, the final movie time point. The number of dorsal and ventral *sox10^+^* cells were counted for the entire z-stack for each individual embryo. To determine the time of the first OPC, the time of the first *sox10^+^* cell to appear in any z-plane was recorded for each embryo. To determine the number of OPC divisions, we counted the number of *sox10^+^* cells that divided across all z-planes for the entirety of the time-lapse movie. Cell divisions were indicated by visualization of an individual *sox10^+^* cell splitting into two independent cells. All quantifications were obtained from a minimum of 10 individual embryos per experimental group.

#### Statistical Analyses

All data analysis, statistics, and plotting were performed in GraphPad Prism (version 11). *p*-values involving only two groups were calculated using an unpaired two-tailed *t*-test. Significance levels were determined using a confidence interval of 95%. The data in the plots and in the text are presented as means ± SEM. A power analysis was performed in R using the “pwr” package with a significance of 0.05, power of 0.8, and standard deviation of 0.35 and the *n* was determined to be 8.76. A minimum of 10 individual embryos per experimental group were used in all quantified studies.

### Data Accessibility

Our single-cell multi-omics dataset can be found at GEO: GSE316142 (Arena et al., 2026).

## Author Contributions

K.A.A and B.H.A designed research, K.A.A performed research, K.A.A analyzed data, K.A.A and B.H.A. wrote the paper, K.A.A, S.J.F, C.G.S, and B.H.A. funded the studies.

## Acknowledgements

This work was supported by the National Institutes of Health (NIH)/National Institute of Neurological Disorders and Stroke (R01 NS124166 to S.J.F., B.H.A., and C.G.S.; R35 NS122191 to B.H.A; and F32 NS134612 to K.A.A). We thank Appel Lab members, past and present, for valuable conversations throughout the course of this project. We thank Rebecca O’Rourke for technical advice for bioinformatics analyses, our zebrafish facility staff for animal care, and Dr. Victor Ruthig and the Developmental Biology Microscopy Consortium for imaging assistance.

The authors declare no conflict of interest.

## KEY RESOURCES

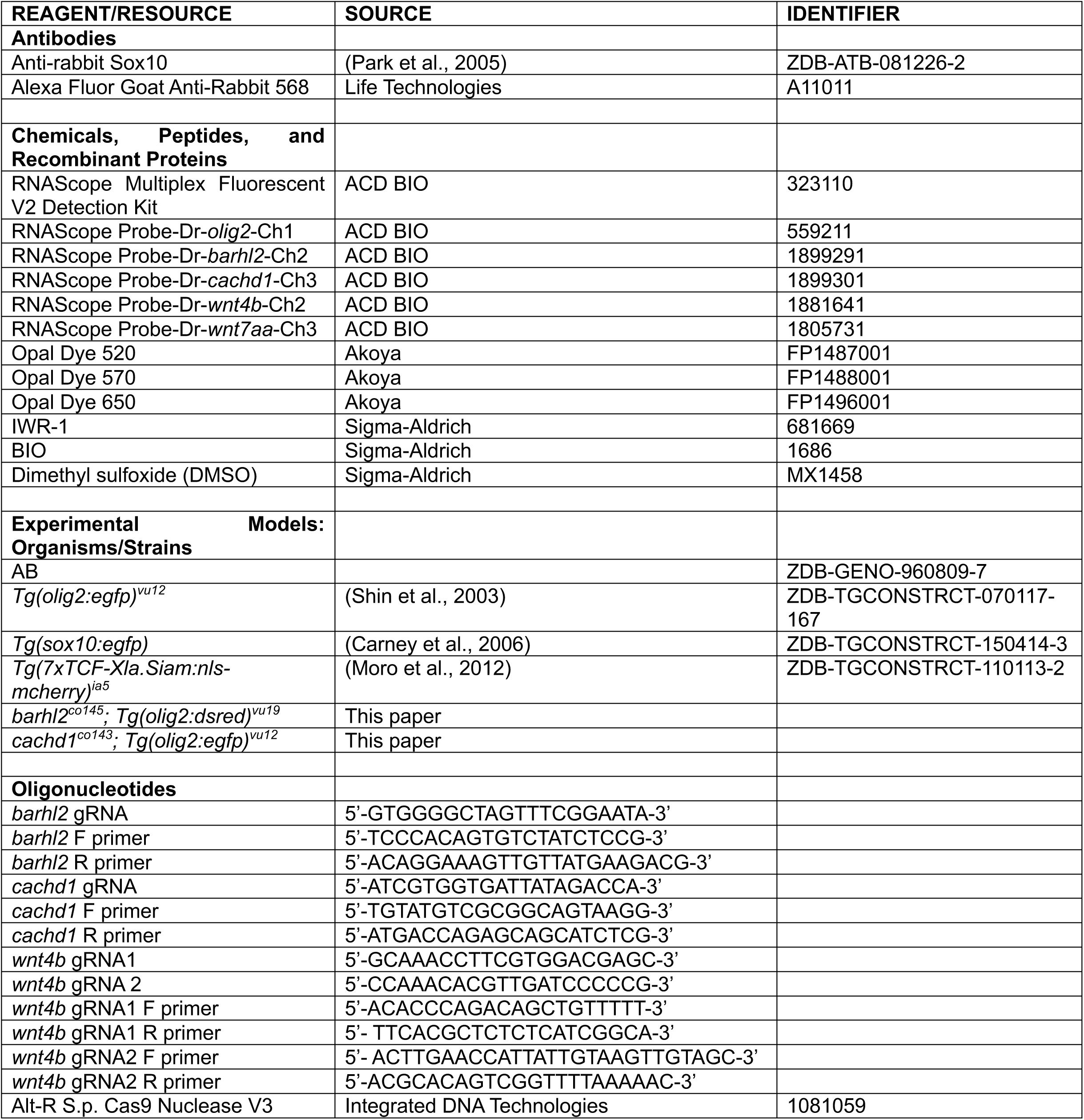

## REFERENCES

Ablinger, C., Eibl, C., Geisler, S. M., Campiglio, M., Stephens, G. J., Missler, M., & Obermair, G. J. (2022). α2δ-4 and Cachd1 Proteins Are Regulators of Presynaptic Functions. International Journal of Molecular Sciences *2022, Vol. 23, Page* 9885, 23(17), 9885. 10.3390/IJMS23179885

Agalliu, D., Takada, S., Agalliu, I., McMahon, A. P., & Jessell, T. M. (2009). Motor Neurons with Axial Muscle Projections Specified by Wnt4/5 Signaling. Neuron, 61(5), 708–720. 10.1016/j.neuron.2008.12.026

Arena, K. A., Kearns, C. A., Ahmed, M., O’Rourke, R., Sagerström, C. G., Franco, S. J., & Appel, B. (2026). Gsx2 regulates oligodendrocyte precursor formation in the zebrafish spinal cord. Developmental Biology, 531, 30–44. 10.1016/J.YDBIO.2026.01.001

Baroti, T., Zimmermann, Y., Schillinger, A., Liu, L., Lommes, P., Wegner, M., & Stolt, C. C. (2016). Transcription factors Sox5 and Sox6 exert direct and indirect influences on oligodendroglial migration in spinal cord and forebrain. GLIA, 64(1), 122–138. 10.1002/GLIA.22919

Bou-Rouphael, J., Doulazmi, M., Eschstruth, A., Abdou, A., & Durand, B. C. (2024). Cerebellar granular neuron progenitors exit their germinative niche via BarH-like1 activity mediated partly by inhibition of T-cell factor. Development (Cambridge), 151(13). 10.1242/DEV.202234

Bou-Rouphael, J., & Durand, B. C. (2021). T-Cell Factors as Transcriptional Inhibitors: Activities and Regulations in Vertebrate Head Development. Frontiers in Cell and Developmental Biology, 9, 784998. 10.3389/FCELL.2021.784998/FULL

Brafman, D., & Willert, K. (2017). Wnt/β-catenin signaling during early vertebrate neural development. Developmental Neurobiology, 77(11), 1239. 10.1002/DNEU.22517

Carney, T. J., Dutton, K. A., Greenhill, E., Delfino-Machín, M., Dufourcq, P., Blader, P., & Kelsh, R. N. (2006). A direct role for Sox10 in specification of neural crest-derived sensory neurons. Development, 133(23), 4619– 4630. 10.1242/DEV.02668

Carr, L., Parkinson, D. B., & Dun, X. P. (2017). Expression patterns of Slit and Robo family members in adult mouse spinal cord and peripheral nervous system. PLoS ONE, 12(2), e0172736. 10.1371/JOURNAL.PONE.0172736

Chen, B., Dodge, M. E., Tang, W., Lu, J., Ma, Z., Fan, C. W., Wei, S., Hao, W., Kilgore, J., Williams, N. S., Roth, M. G., Amatruda, J. F., Chen, C., & Lum, L. (2009). Small molecule-mediated disruption of Wnt-dependent signaling in tissue regeneration and cancer. Nature Chemical Biology, 5(2), 100. 10.1038/NCHEMBIO.137

Costantino, I., Meng, A., & Ravits, J. (2024). Alternatively spliced ELAVL3 cryptic exon 4a causes ELAVL3 downregulation in ALS TDP-43 proteinopathy. Acta Neuropathologica, 147(1), 93. 10.1007/S00401-024-02732-Y

Dai, Z. M., Sun, S., Wang, C., Huang, H., Hu, X., Zhang, Z., Lu, Q. R., & Qiu, M. (2014). Stage-Specific Regulation of Oligodendrocyte Development by Wnt/β-Catenin Signaling. Journal of Neuroscience, 34(25), 8467–8473. 10.1523/JNEUROSCI.0311-14.2014

Fancy, S. P. J., Baranzini, S. E., Zhao, C., Yuk, D. I., Irvine, K. A., Kaing, S., Sanai, N., Franklin, R. J. M., & Rowitch, D. H. (2009). Dysregulation of the Wnt pathway inhibits timely myelination and remyelination in the mammalian CNS. Genes & Development, 23(13), 1571. 10.1101/GAD.1806309

Floriddia, E. M., Lourenço, T., Zhang, S., van Bruggen, D., Hilscher, M. M., Kukanja, P., Gonçalves dos Santos, J. P., Altınkök, M., Yokota, C., Llorens-Bobadilla, E., Mulinyawe, S. B., Grãos, M., Sun, L. O., Frisén, J., Nilsson, M., & Castelo-Branco, G. (2020). Distinct oligodendrocyte populations have spatial preference and different responses to spinal cord injury. Nature Communications *2020 11:1*, 11(1), 1–15. 10.1038/s41467-020-19453-x

Freese, J. L., Pino, D., & Pleasure, S. J. (2009). Wnt signaling in development and disease. Neurobiology of Disease, 38(2), 148. 10.1016/j.nbd.2009.09.003

Fu, H., Kesari, S., & Cai, J. (2012). Tcf7l2 is Tightly Controlled During Myelin Formation. Cellular and Molecular Neurobiology, 32(3), 345. 10.1007/S10571-011-9778-Y

Guo, F., Lang, J., Sohn, J., Hammond, E., Chang, M., & Pleasure, D. (2015). Canonical Wnt signaling in the oligodendroglial lineage-puzzles remain. GLIA, 63(10), 1671–1693. 10.1002/glia.22813

Guo, F., & Wang, Y. (2023). TCF7l2, a nuclear marker that labels premyelinating oligodendrocytes and promotes oligodendroglial lineage progression. Glia, 71(2), 143–154. 10.1002/GLIA.24249

Hao, Y., Stuart, T., Kowalski, M. H., Choudhary, S., Hoffman, P., Hartman, A., Srivastava, A., Molla, G., Madad, S., Fernandez-Granda, C., & Satija, R. (2023). Dictionary learning for integrative, multimodal and scalable single-cell analysis. Nature Biotechnology *2023 42:2*, 42(2), 293–304. 10.1038/s41587-023-01767-y

Inohaya, K., Takano, Y., & Kudo, A. (2010). Production of Wnt4b by floor plate cells is essential for the segmental patterning of the vertebral column in medaka. Development, 137(11), 1807–1813. 10.1242/dev.051540

Jiang, M., Yu, D., Xie, B., Huang, H., Lu, W., Qiu, M., & Dai, Z. M. (2020). WNT signaling suppresses oligodendrogenesis via Ngn2-dependent direct inhibition of Olig2 expression. Molecular Brain *2020 13:1,* 13(1), 155-. 10.1186/S13041-020-00696-0

Juraver-Geslin, H. A., Ausseil, J. J., Wassef, M., & Durand, B. C. (2011). Barhl2 limits growth of the diencephalic primordium through Caspase3 inhibition of β-catenin activation. Proceedings of the National Academy of Sciences of the United States of America, 108(6), 2288–2293. 10.1073/pnas.1014017108

Kimmel, C. B., Ballard, W. W., Kimmel, S. R., Ullmann, B., & Schilling, T. F. (1995). Stages of embryonic development of the zebrafish. Developmental Dynamics, 203(3), 253–310. 10.1002/aja.1002030302

Kucenas, S., Snell, H., & Appel, B. (2008). Nkx2.2a promotes specification and differentiation of a myelinating subset of oligodendrocyte lineage cells in zebrafish. Neuron Glia Biology, 4(2), 71–81. 10.1017/S1740925X09990123

Langseth, A. J., Munji, R. N., Choe, Y., Huynh, T., Pozniak, C. D., & Pleasure, S. J. (2010). Wnts Influence the Timing and Efficiency of Oligodendrocyte Precursor Cell Generation in the Telencephalon. The Journal of Neuroscience, 30(40), 13367. 10.1523/JNEUROSCI.1934-10.2010

Liang, X., Song, M. R., Xu, Z. G., Lanuza, G. M., Liu, Y., Zhuang, T., Chen, Y., Pfaff, S. L., Evans, S. M., & Sun, Y. (2011). Isl1 Is required for multiple aspects of motor neuron development. Molecular and Cellular Neurosciences, 47(3), 215. 10.1016/J.MCN.2011.04.007

Liu, A., Majumdar, A., Schauerte, H. E., Haffter, P., & Drummond, I. A. (2000). Zebrafish wnt4b expression in the floor plate is altered in sonic hedgehog and gli-2 mutants. Mechanisms of Development, 91(1–2), 409–413. 10.1016/S0925-4773(99)00308-1

Masahira, N., Takebayashi, H., Ono, K., Watanabe, K., Ding, L., Furusho, M., Ogawa, Y., Nabeshima, Y. ichi, Alvarez-Buylla, A., Shimizu, K., & Ikenaka, K. (2006). Olig2-positive progenitors in the embryonic spinal cord give rise not only to motoneurons and oligodendrocytes, but also to a subset of astrocytes and ependymal cells. Developmental Biology. 10.1016/j.ydbio.2006.02.029

Megason, S. G., & McMahon, A. P. (2002). A mitogen gradient of dorsal midline Wnts organizes growth in the CNS. Development (Cambridge, England), 129(9), 2087–2098. 10.1242/DEV.129.9.2087

Meijer, L., Skaltsounis, A. L., Magiatis, P., Polychronopoulos, P., Knockaert, M., Leost, M., Ryan, X. P., Vonica, C. A., Brivanlou, A., Dajani, R., Crovace, C., Tarricone, C., Musacchio, A., Roe, S. M., Pearl, L., & Greengard, P. (2003). GSK-3-Selective Inhibitors Derived from Tyrian Purple Indirubins. Chemistry & Biology, 10(12), 1255–1266. 10.1016/J.CHEMBIOL.2003.11.010

Moro, E., Ozhan-Kizil, G., Mongera, A., Beis, D., Wierzbicki, C., Young, R. M., Bournele, D., Domenichini, A., Valdivia, L. E., Lum, L., Chen, C., Amatruda, J. F., Tiso, N., Weidinger, G., & Argenton, F. (2012). In vivo Wnt signaling tracing through a transgenic biosensor fish reveals novel activity domains. Developmental Biology, 366(2), 327–340. 10.1016/J.YDBIO.2012.03.023

Muroyama, Y., Fujihara, M., Ikeya, M., Kondoh, H., & Takada, S. (2002). Wnt signaling plays an essential role in neuronal specification of the dorsal spinal cord. Genes & Development, 16(5), 548–553. 10.1101/GAD.937102

Nery, S., Wichterle, H., & Fishell, G. (2001). Sonic hedgehog contributes to oligodendrocyte specification in the mammalian forebrain. Development (Cambridge, England), 128(4), 527–540. 10.1242/DEV.128.4.527

Oh, S., Huang, X., & Chiang, C. (2005). Specific requirements of sonic hedgehog signaling during oligodendrocyte development. Developmental Dynamics, 234(3), 489–496. 10.1002/DVDY.20422

Oh, S. O., Huang, X., Liu, J., Litingtung, Y., & Chiang, C. (2009). Shh and Gli3 activities are required for timely generation of motor neuron progenitors. Developmental Biology, 331(2), 261–269. 10.1016/J.YDBIO.2009.05.539

Ovek Baydar, D., Rauluseviciute, I., Aronsen, D. R., Blanc-Mathieu, R., Bonthuis, I., De Beukelaer, H., Ferenc, K., Jegou, A., Kumar, V., Lemma, R. B., Lucas, J., Pochon, M., Yun, C. M., Ramalingam, V., Deshpande, S. S., Patel, A., Marinov, G. K., Wang, A. T., Aguirre, A., … Mathelier, A. (2025). JASPAR 2026: expansion of transcription factor binding profiles and integration of deep learning models. Nucleic Acids Research, 54(D1), D184. 10.1093/NAR/GKAF1209

Park, H. C., Boyce, J., Shin, J., & Appel, B. (2005). Oligodendrocyte Specification in Zebrafish Requires Notch-Regulated Cyclin-Dependent Kinase Inhibitor Function. Journal of Neuroscience, 25(29), 6836–6844. 10.1523/JNEUROSCI.0981-05.2005

Park, H. C., Mehta, A., Richardson, J. S., & Appel, B. (2002). olig2 is required for zebrafish primary motor neuron and oligodendrocyte development. Developmental Biology. 10.1006/dbio.2002.0738

Park, H. C., Shin, J., & Appel, B. (2004). Spatial and temporal regulation of ventral spinal cord precursor specification by Hedgehog signaling. Development, 131(23), 5959–5969. 10.1242/DEV.01456

Powell, G. T., Faro, A., Zhao, Y., Stickney, H., Novellasdemunt, L., Henriques, P., Gestri, G., White, E. R., Ren, J., Lu, W., Young, R. M., Hawkins, T. A., Cavodeassi, F., Schwarz, Q., Dreosti, E., Raible, D. W., Li, V. S. W., Wright, G. J., Jones, E. Y., & Wilson, S. W. (2024). Cachd1 interacts with Wnt receptors and regulates neuronal asymmetry in the zebrafish brain. Science (New York, N.Y.), 384(6695), 573. 10.1126/SCIENCE.ADE6970

Ravanelli, A. M., & Appel, B. (2015). Motor neurons and oligodendrocytes arise from distinct cell lineages by progenitor recruitment. Genes & Development, 29(23), 2504. 10.1101/GAD.271312.115

Ravanelli, A. M., Kearns, C. A., Powers, R. K., Wang, Y., Hines, J. H., Donaldson, M. J., & Appel, B. (2018). Sequential specification of oligodendrocyte lineage cells by distinct levels of Hedgehog and Notch signaling. Developmental Biology, 444(2), 93–106. 10.1016/J.YDBIO.2018.10.004

Scala, M., Khan, K., Beneteau, C., Fox, R. G., von Hardenberg, S., Khan, A., Joubert, M., Fievet, L., Musquer, M., Le Vaillant, C., Holsclaw, J. K., Lim, D., Berking, A. C., Accogli, A., Giacomini, T., Nobili, L., Striano, P., Zara, F., Torella, A., … Wells, M. F. (2023). Biallelic loss-of-function variants in CACHD1 cause a novel neurodevelopmental syndrome with facial dysmorphism and multisystem congenital abnormalities. Genetics in Medicine : Official Journal of the American College of Medical Genetics, 26(4), 101057. 10.1016/J.GIM.2023.101057

Schubert, M., Klinger, B., Klünemann, M., Sieber, A., Uhlitz, F., Sauer, S., Garnett, M. J., Blüthgen, N., & Saez-Rodriguez, J. (2018). Perturbation-response genes reveal signaling footprints in cancer gene expression. Nature Communications *2017 9:1*, 9(1), 20. 10.1038/s41467-017-02391-6

Scott, K., O’Rourke, R., Winkler, C. C., Kearns, C. A., & Appel, B. (2021). Temporal single-cell transcriptomes of zebrafish spinal cord pMN progenitors reveal distinct neuronal and glial progenitor populations. Developmental Biology, 479, 37–50. 10.1016/J.YDBIO.2021.07.010

Sena, E., Rocques, N., Borday, C., Amin, H. S. M., Parain, K., Sitbon, D., Chesneau, A., & Duran, B. C. (2019). Barhl2 maintains t cell factors as repressors and thereby switches off the wnt/β-catenin response driving spemann organizer formation. Development (Cambridge), 146(10). 10.1242/DEV.173112/48830

Shin, J., Park, H. C., Topczewska, J. M., Madwsley, D. J., & Appel, B. (2003). Neural cell fate analysis in zebrafish using olig2 BAC transgenics. Methods in Cell Science, 25(1–2), 7–14. 10.1023/B:MICS.0000006847.09037.3A

Soomro, S. H., Jie, J., & Fu, H. (2018). Oligodendrocytes Development and Wnt Signaling Pathway. International Journal of Human Anatomy, 1(3), 17. 10.14302/ISSN.2577-2279.IJHA-18-2407

Stuart, T., Srivastava, A., Madad, S., Lareau, C. A., & Satija, R. (2021). Single-cell chromatin state analysis with Signac. Nature Methods *2021 18:11,* 18(11), 1333–1341. 10.1038/s41592-021-01282-5

Tawk, M., Makoukji, J., Belle, M., Fonte, C., Trousson, A., Hawkins, T., Li, H., Ghandour, S., Schumacher, M., & Massaad, C. (2011). Wnt/β-Catenin Signaling Is an Essential and Direct Driver of Myelin Gene Expression and Myelinogenesis. The Journal of Neuroscience, 31(10), 3729. 10.1523/JNEUROSCI.4270-10.2011

Tekki-Kessaris, N., Woodruff, R., Hall, A. C., Gaffield, W., Kimura, S., Stiles, C. D., Rowitch, D. H., & Richardson, W. D. (2001). Hedgehog-dependent oligodendrocyte lineage specification in the telencephalon. Development (Cambridge, England), 128(13), 2545–2554. 10.1242/DEV.128.13.2545

Tyurina, O. V., Guner, B., Popova, E., Feng, J., Schier, A. F., Kohtz, J. D., & Karlstrom, R. O. (2005). Zebrafish Gli3 functions as both an activator and a repressor in Hedgehog signaling. Developmental Biology, 277(2), 537–556. 10.1016/J.YDBIO.2004.10.003

Ulloa, F., Itasaki, N., & Briscoe, J. (2007). Inhibitory Gli3 Activity Negatively Regulates Wnt/β-Catenin Signaling. Current Biology, 17(6), 545–550. 10.1016/j.cub.2007.01.062

Wang, H., & Matise, M. P. (2016). Tcf7l2/Tcf4 Transcriptional Repressor Function Requires HDAC Activity in the Developing Vertebrate CNS. PLOS ONE, 11(9), e0163267. 10.1371/JOURNAL.PONE.0163267

Wang, L. C., & Almazan, G. (2016). Role of Sonic Hedgehog Signaling in Oligodendrocyte Differentiation. Neurochemical Research, 41(12), 3289–3299. 10.1007/S11064-016-2061-3

Warf, B. C., Fok-Seang, J., & Miller, R. H. (1991). Evidence for the ventral origin of oligodendrocyte precursors in the rat spinal cord. The Journal of Neuroscience : The Official Journal of the Society for Neuroscience, 11(8), 2477–2488. 10.1523/JNEUROSCI.11-08-02477.1991

Xiao, G., Du, J., Wu, H., Ge, X., Xu, X., Yang, A., Zhu, Y., Hu, X., Zheng, K., Zhu, Q., & Qiu, M. (2019). Differential Inhibition of Sox10 Functions by Notch-Hes Pathway. Cellular and Molecular Neurobiology *2019 40:4*, 40(4), 653–662. 10.1007/S10571-019-00764-7

Yu, W., McDonnell, K., Taketo, M. M., & Bai, C. B. (2008). Wnt signaling determines ventral spinal cord cell fates in a time-dependent manner. Development, 135(22), 3687–3696. 10.1242/DEV.021899

Zhang, S., Wang, Y., Zhu, X., Song, L., Zhan, X., Ma, E., McDonough, J., Fu, H., Cambi, F., Grinspan, J., & Guo, F. (2021). The Wnt Effector TCF7l2 Promotes Oligodendroglial Differentiation by Repressing Autocrine BMP4-Mediated Signaling. The Journal of Neuroscience : The Official Journal of the Society for Neuroscience, 41(8), 1650–1664. 10.1523/JNEUROSCI.2386-20.2021

Zhao, C., Deng, Y., Liu, L., Yu, K., Zhang, L., Wang, H., He, X., Wang, J., Lu, C., Wu, L. N., Weng, Q., Mao, M., Li, J., Van Es, J. H., Xin, M., Parry, L., Goldman, S. A., Clevers, H., & Lu, Q. R. (2016). Dual regulatory switch through interactions of Tcf7l2/Tcf4 with stage-specific partners propels oligodendroglial maturation. Nature Communications *2016 7:1*, 7(1), 1–15. 10.1038/ncomms10883

Zhou, Q., Choi, G., & Anderson, D. J. (2001). The bHLH transcription factor Olig2 promotes oligodendrocyte differentiation in collaboration with Nkx2.2. Neuron, 31(5), 791–807. 10.1016/S0896-6273(01)00414-7

